# Mouse eradication is required to prevent local extinction of an endangered seabird on an oceanic island

**DOI:** 10.1101/2020.11.16.385997

**Authors:** Christopher W. Jones, Michelle M. Risi, Alexis M. Osborne, Peter G. Ryan, Steffen Oppel

**Author notes:** Corresponding Author, Twitter; @SteffOpp.

## Abstract

Petrels (Procellariidae) are a highly diverse family of seabirds, many of which are globally threatened due to the impact of invasive species on breeding populations. While predation by invasive cats and rats has led to the extinction of petrel populations, the impact of invasive house mice *Mus musculus* is slower and less well documented. However, mice impact small burrow-nesting species such as MacGillivray’s prion *Pachyptila macgillivrayi,* a species classified as endangered because it has been extirpated on islands in the Indian Ocean by introduced rodents. We use historic abundance data and demographic monitoring data from 2014 to 2020 to predict the population trajectory of MacGillivray’s prion on Gough Island with and without a mouse eradication using a stochastic integrated population model. Given very low annual breeding success (0.01 fledglings per breeding pair in ‘poor’ years (83%) or 0.38 in ‘good’ years (17%), n = 320 nests over 6 years) mainly due to mouse predation, our model predicted that the population collapsed from ~3.5 million pairs in 1956 to an estimated 175,000 pairs in 2020 despite reasonably high adult survival probability *(ϕ* = 0.901). Based on these parameters, the population is predicted to decline at a rate of 9% per year over the next 36 years without a mouse eradication, with a 31% probability that by 2057 the MacGillivray’ prion population would become extremely vulnerable to extinction. Our models predict population stability (λ = 1.01) and a lower extinction risk (<10%) if mouse eradication on Gough Island restores annual breeding success to 0.519, which is in line with that of closely-related species on predator-free islands. This study demonstrates the devastating impacts that introduced house mice can have on small burrowing petrels and highlights the urgency to eradicate invasive mammals from oceanic islands.

## Introduction

In the Southern Ocean, petrels (Procellariidae) are the most diverse, widespread and abundant seabird group. These birds have long lifespans, low fecundity and take several years to reach sexual maturity; and these traits have likely evolved because they mostly breed on remote oceanic islands in the absence of mammalian predators (Warham 1990; Brooke 2004). The introduction of invasive mammals on oceanic islands has negatively affected petrel populations by reducing breeding success mainly through the predation of chicks and eggs (Brooke et al. 2018), and occasionally through the predation of adult birds (Van Rensburg and Bester 1988; Keitt et al. 2002; Rayner et al. 2007; Faulquier et al. 2009). Consequently, about 50% of extant petrel species are currently classified as globally threatened (Dias et al. 2019).

The decline of many petrel populations has largely been attributed to the introduction of rats *Rattus* spp. and feral cats *Felis catus* to their breeding islands (Brooke et al. 2018). However, house mice *Mus musculus* can have significant impacts on seabirds at islands where they are the only introduced predator, such as Gough Island in the central South Atlantic Ocean or Marion Island in the South-Western Indian Ocean (Cuthbert and Hilton 2004; Wanless et al. 2007; Wanless et al. 2009; Cuthbert et al. 2013; Davies et al. 2015; Dilley et al. 2015; Dilley et al. 2016; Caravaggi et al. 2018; Dilley et al. 2018; Jones et al. 2019). While rats and cats are typical predators that start depredating seabird eggs and chicks soon after the invasion of an island, this behaviour appears to evolve gradually in invasive mouse populations only at a later stage of invasion when other primarily exploited food sources have been depleted (McClelland et al. 2018; Russell et al. 2020).

Prions *(Pachyptila* spp.) are small (130–250 g), planktivorous petrels which breed in large numbers on islands throughout the Southern Ocean (Brooke 2004). Their taxonomy is still debated, and until recently, McGillivray’s prion *P. macgillivrayi* was considered to be a subspecies of Salvin’s prion *P. salvini* or the Indian Ocean counterpart of the broad-billed prion *P. vittata* (Bretagnolle et al. 1990; Warham 1990). Consequently, only the Broad-billed Prion *Pachyptila vittata* was thought to breed at Tristan da Cunha and Gough Island. However, morphological, ecological and genetic evidence now indicates that MacGillivray’s prion breeds sympatrically with *P. vittata* on Gough Island (Ryan et al. 2014; Birdlife International 2020; Jones et al. 2020). The only other population of MacGillivray’s prion is a tiny relict population on a rock stack near St Paul Island in the central Indian Ocean (Worthy and Jouventin 1999; Jiguet et al. 2007), and MacGillivray’s prions therefore have a disjunct distribution in both the South Atlantic and the Southern Indian Ocean (Birdlife-International 2020).

Despite the geographic distribution, MacGillivray’s prion is nearly endemic to Gough Island. The species was once abundant on Amsterdam and St Paul islands, but was extirpated from both islands by introduced mammalian predators (Worthy and Jouventin 1999; Jiguet et al. 2007), and the Southern Indian Ocean population of the species is now very small, with an estimated 150-200 pairs mostly confined to La Roche Quille, a rock stack 150 m off the coast of St Paul Island (Tollu 1984; Jiguet et al. 2007). By contrast, hundreds of thousands of prions nest on Gough, but the population size and trends are poorly understood as these burrow-nesting and nocturnal birds are exceptionally difficult to census. Swales (1965) estimated that there were approximately 10 million pairs of prions breeding on Gough Island in 1955/56, and Cuthbert (2004) estimated 1.5-2.0 million pairs of prions in 2000/01, but both estimates are surrounded by large uncertainty. The population size of MacGillivray’s prion – a subset of these combined estimates of all prions – is therefore equally uncertain. Based on estimates of their proportions in skua middens collected around the island, MacGillivray’s prions comprise 40-50% of prions breeding on Gough Island (Jones 2018), which suggests that there were approximately 600 000 – 1 000 000 pairs of MacGillivray’s prions in 2000/01, and thus > 99% of the global population.

Given the extinction of MacGillivray’s prion on Amsterdam and St Paul islands due to invasive mammalian predators, there is a justified concern that the species may suffer a similar fate on Gough, but demographic data to estimate extinction probability are lacking so far. Here we provide an updated assessment of the population status of the MacGillivray’s prion population breeding at Gough Island by projecting the latest estimate three generations into the future using a stochastic integrated population model. We estimate demographic parameters from a monitored colony of McGillivray’s prions on Gough Island, and examine whether these parameters are consistent with the decline between the only two population assessments in 1956 and 2001. Finally, we explore what effect a planned mouse eradication will have on the population assuming that breeding success would be restored to levels of the same species breeding on an island free of introduced predators (Jiguet et al. 2007).

## Methods

### Study area and focal species

Gough is a volcanic island (65.8 km^2^; 40°21′S, 9·53′W, Fig. 1) located approximately 2500 km southwest of South Africa, and 400 km southeast of Tristan da Cunha. Gough has steep mountainous terrain rising to ~910 m above sea level (asl), and is inhabited by 23 breeding seabird species, including the two sympatrically breeding prion species. Both prions nest in caves and burrows from sea level to 700 m asl. Breeding is highly synchronous within these prion species, but MacGillivray’s prions breed approximately 3 months after broad-billed prions, and lay eggs from late November to early December and fledge chicks in February– March (Jones et al. 2020).

**Figure 1:**
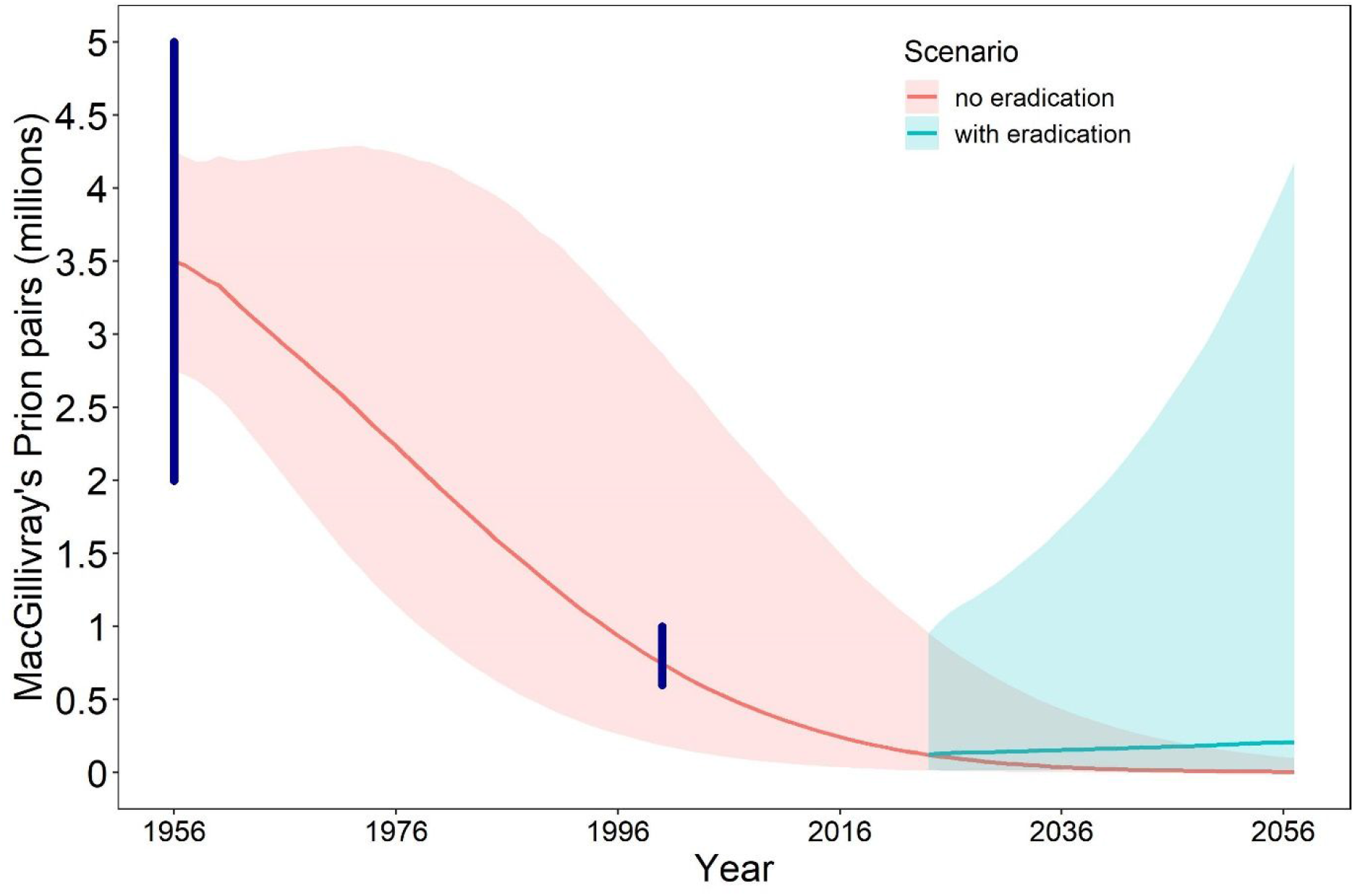
Extrapolated population trajectory of the number of MacGillivray’s prion breeding pairs between 1956 and 2057 on Gough Island (with 50% credible interval) based on a population model informed by nest monitoring and mark-recapture data collected between 2014 and 2019 with and without a mouse eradication planned in 2021. The dark blue vertical lines indicate the range of abundance estimated from field surveys in 1956 and 2000.

The conservation status of MacGillivray’s prion was first assessed for the IUCN Red List in 2016, because it was not recognised as a species by BirdLife International previously. MacGillivray’s prion is classified as Endangered under Criterion B2a (two populations with a total breeding area <70 km^2^) and B2b because the species has experienced very poor breeding success (< 10%) at Gough Island in recent years due to high chick mortality caused by mouse predation (Dilley et al. 2015; Caravaggi et al. 2018; Birdlife-International 2020).

### Demographic monitoring

A monitoring colony of MacGillivray’s prions was set up in 2014 in a small accessible cave (known as ‘Prion Cave’), which is approximately 30 m long and is typically occupied by 100 – 200 accessible MacGillivray’s prions. This cave offers an opportunity to study a species that typically nests in inaccessible small caves and jumbled boulder crevasses, and our study cave is unlikely to offer systematic survival or breeding success advantages or disadvantages to prions: mouse predation on burrowing petrel chicks is widespread within and outside the cave (Dilley *et al.,* 2015; Caravaggi *et al.,* 2018), and skua predation of young or adult birds can similarly occur for birds outside large or small caves, crevasses or burrows. We therefore assume that the demographic parameters measured in this cave colony are representative of the prion population on Gough Island.

We collected data on breeding success and adult survival at this site each year from 2014 – 2020. At the peak of the egg laying period during the last week of November, 50 – 70 nests were individually marked and these nests were checked weekly throughout each season until chicks fledged in late February/early March. Because prions lay only one egg per season, breeding success was defined as the proportion of nests that successfully fledged a chick. Breeding adults were ringed at their nests with uniquely numbered metal rings. These metal ring numbers were recorded every year for all breeders during incubation.

Some birds in this cave were ringed without belonging to a monitored breeding pair, and may have transited the breeding area to inaccessible parts of the cave, where prions also breed, but cannot be monitored. These transient individuals may have had a lower probability to be recaptured, and we therefore accounted for the presence of such individuals in our survival model (Genovart and Pradel 2019).

### Population model

Removing invasive species from islands can have enormous benefits for seabirds (Jones et al. 2016; Holmes et al. 2019), mostly because of increased breeding success (Lavers et al. 2010; Brooke et al. 2018). Because mouse predation causes low breeding success of prions on Gough Island, we explored whether mouse eradication could reverse assumed population declines.

We used a stochastic integrated population model to estimate demographic parameters and population size for Gough Island while accounting for all environmental and parameter uncertainty (Schaub and Abadi 2011; Zipkin and Saunders 2018; Plard et al. 2019). Briefly, this model was based on an age-structured matrix model with four different age classes, adapted from a similar model for MacGillivray’s prion in the Indian Ocean (Jiguet et al. 2007). We introduced further flexibility to relax several assumptions and ensure that all parameter uncertainty was incorporated into population projections to avoid overconfident predictions. Our model assumed an equal sex ratio at hatching and that 85 – 95% of all adult birds (4 years and older) would attempt to reproduce every year (Brooke et al. 2010). The number of birds in each age class was randomly drawn for each year based on the abundance and survival probability of birds from the previous year (Abadi et al. 2010), and the productivity of adult breeding birds. To estimate productivity, we used the annual number of fledglings corrected for the annual number of observed pairs with a Poisson error distribution, and we assumed that this productivity followed a bimodal distribution, with ‘good’ years and ‘poor’ years. We estimated the annual productivity for ‘good’ and ‘poor’ years separately, and the frequency at which ‘good’ years occurred.

We estimated annual adult survival probability *ϕ* with a Cormack-Jolly-Seber model using the recapture history of individually marked birds from 2014 – 2019. We first explored the most parsimonious temporal parameterisation of the survival model, and assessed the goodness-of-fit of the model for our data set. We fitted four competing models that explored that annual survival and recapture probability either varied by year or were constant, and selected the most parsimonious of these four models as indicated by the lowest Akaike Information Criterion adjusted for small sample size (Burnham and Anderson 2002). This preliminary model selection indicated that a model formulation with constant annual survival and annually varying recapture probability was the most parsimonious (Table S1), and we therefore adopted this parameterisation. We then fitted this model either with all individuals, or with just individuals that were captured more than once, to remove the effect of transients (Pradel et al. 1997). For each model we assessed goodness-of-fit using a posterior predictive check of the discrepancy between expected and observed data (Kéry and Schaub 2012; Conn et al. 2018), which indicated that a model excluding transients was inappropriate (Bayesian p-value = 0.006), but that including all birds resulted in an adequate model fit (Bayesian p-value = 0.42, Fig. S1). To appropriately account for the presence of transient individuals in our data set, we therefore adopted a multi-event model parameterisation that allowed for emigration in the transition probability between subsequent encounter occasions for each individual (Genovart and Pradel 2019).

Because insufficient data existed for birds marked as chicks, we were unable to empirically estimate the annual survival probability of juveniles. We assumed that birds would start breeding at the age of four years, and that immature survival after the first year would be identical to adult survival (Jiguet et al. 2007). Given that extremely few empirical data for the juvenile survival of small petrel species exist, we followed Jiguet et al. (2007) and adopted the juvenile survival of Balearic Shearwaters *(Puffinus mauretanicus)* (Oro et al. 2004). We specified juvenile survival as a randomly drawn variable between the values of 0.63 and 0.78, reflecting the entire 95% confidence interval of empirical data (Oro et al. 2004), and the values used in similar population models for petrels (Jiguet et al. 2007; Brooke et al. 2010).

To constrain population projections from this demographic model to match with the only two empirical estimates of the MacGillivray’s prion population size on Gough Island (1956: 2-5 million pairs; 2000: 600,000 – 1.2 million pairs), we initiated the model in 1956 and drew the original population size from a random uniform distribution between 2 and 5 million breeding pairs. Because mouse predation on seabird chicks has likely increased recently (Ryan 2010, Dilley *et al.* 2015; Jones *et al.* 2019), it was unrealistic to expect that productivity had been constantly low since 1956. We therefore assumed that productivity in 1956 was substantially higher, and specified this by an increasing frequency of ‘good’ breeding years. The frequency of ‘good’ breeding years was then assumed to decrease linearly until our empirical estimates began in 2014, and the prior for the distribution of original frequency of ‘good’ breeding years was chosen to ensure that the population trajectory based on our demographic parameters would lie between 600,000 – 1.2 million pairs in the year 2000.

We used a Bayesian framework for inference and parameter estimation because it provided the required flexibility and allowed for the incorporation of existing information to inform prior distributions for demographic parameters (Wade 2000; Brooks et al. 2004; Schaub et al. 2007). Specifically, we used an informative prior (0.7-1) for annual adult survival and for productivity (0 – 0.5) given what is known about the species from Gough and other populations (Jiguet et al. 2007; Caravaggi et al. 2018). We fitted the population model in JAGS (Plummer 2012) called from R 4.0.1 (R-Core-Team 2019) via the package ‘jagsUI’ (Kellner 2016). We ran three Markov chains each with 150,000 iterations and discarded the first 50,000 iterations. We tested for convergence using the Gelman-Rubin diagnostic and confirmed that R-hat was <1.01 for all parameters. We present posterior estimates of parameters with 95% credible intervals. Code and data to replicate the analysis and extract the output of all 100,000 simulations is available at https://github.com/steffenoppel/MAPR.

### Population projections under different scenarios

Because Gough Island is an extremely remote, rugged, and uninhabited island, very few management options exist to reduce the adverse effects of invasive mice. Long-term control operations or measures to protect individual seabird nests are not practically feasible on Gough at a scale that could be expected to benefit seabird populations. However, the eradication of mice from Gough Island is feasible, and could be completed in a single winter based on similar operations elsewhere (Broome et al. 2019; Horn et al. 2019). We therefore quantified the future population trajectory of MacGillivray’s prions either with or without a successful mouse eradication on Gough Island in 2021. Because a mouse eradication would occur over a short period (3 months) in mid-winter between two successive breeding seasons (McClelland 2019), we assumed that after 2021 productivity of prions would either remain the same as between 2014 and 2020 (no mouse eradication), or increase to a level observed on an island free of introduced predators (with successful mouse eradication: 0.519, (Jiguet et al. 2007). We then projected the MacGillivray’s prion population on Gough 36 years (three generations; Bird et al. 2020) into the future while accounting for uncertainty and temporal stochasticity in demographic parameters. Our projections included both parameter uncertainty and process variance, because future populations were drawn using binomial trials with parameters allowed to vary in each simulation. These projections therefore account for all the uncertainty associated with demographic parameters and environmental stochasticity.

We calculated the future population growth rate as the geometric mean over the annual growth rates calculated as the population in year *t*+1 over the population in year *t*. We present this future population growth rate to assess whether the mouse eradication would stabilize (growth rate ≥ 1) the prion population on Gough. We also present the probability of becoming extremely vulnerable to extinction by 2057, calculated as the proportion of population simulations under both scenarios where the total number of breeding pairs was <250 (Frankham et al. 2014).

## Results

We monitored 320 MacGillivray’s prion nests between 2014 and 2019, and breeding failure was nearly complete in five out of six years, with no fledged chicks in 2014 (n = 63 nests), 2017 (66) and 2019 (50), and one fledgling in 2015 (40) and 2018 (50); 2016 was the only year when there was higher breeding success (37%, n = 51 nests), and the frequency of ‘good’ breeding years was therefore 17%. We ringed 226 adult individuals between 2014 and 2018, and 71 (31%) of these birds were never recorded after initial capture. The remaining 155 adult birds were recorded on average four times (range 1 – 10 recaptures).

Our population model estimated productivity as 0.01 fledglings per breeding pair in ‘poor’ years, and 0.38 in ‘good’ years, with the frequency of ‘good’ years decreasing from 0.91 in 1956 to 0.17 in 2014 (Table 1). Median annual adult survival probability was estimated as 0.901, but with considerable uncertainty due to the presence of transient individuals (Table 1).

**Table 1.** Demographic parameters for MacGillivray’s prion on Gough Island (median and lower and upper 95% credible limits) used in a population model based on nest monitoring and mark-recapture data collected between 2014 and 2019. Productivity represents chicks fledged per pair, and varies between ‘good’ and ‘poor’ years. The original frequency of ‘good’ breeding years and the decline rate of this frequency were estimated by constraining the model to be consistent with available total population estimates in 1956 and 2000 (Fig. 1). Population growth rate was estimated for 2021 – 2057 assuming two scenarios of future productivity, with or without mouse eradication in 2021.

| Parameter | Median (95% credible interval) |
| --- | --- |
| productivity (poor year) | 0.01 (0.002 – 0.027) |
| productivity (good year) | 0.386 (0.238 – 0.581) |
| original frequency of good breeding year (1956)* | 0.91 (0.881 – 0.939) |
| annual decline in frequency of good breeding year* | -0.013 (-0.015 – -0.011) |
| current frequency of good breeding year (2014-2019) | 0.169 (0.029 – 0.292) |
| first year survival probability* | 0.705 (0.634 – 0.776) |
| annual adult survival probability | 0.901 (0.825 – 0.98) |
| annual population growth rate (no eradication) | 0.908 (0.859 – 0.986) |
| annual population growth rate (with eradication) | 1.01 (0.929 – 1.097) |

\* Note that these parameters reflect prior distributions and the constraint on population estimates in 1956 and 2000, but were not estimated from empirical data.

Based on these demographic parameters we projected the MacGillivray’s prion population on Gough to decrease at a rate of 9% per year over the next 36 years without mouse eradication (Table 1). Using this projection from the last population assessment in 2000, the MacGillivray’s prion population on Gough was estimated at 175,000 pairs (95% credible interval 24,000 – 1,210,000) in 2020, which represents ~5% of the population size in 1956 (Fig. 1). Assuming a successful mouse eradication in 2021, and an increase in productivity to 0.519 fledglings per breeding pair, the future population growth rate would likely be stable (mean 1.01, Table 1). However, we caution that given the large uncertainty in demographic parameters (fully accounted for in our projections), 8% of our 100,000 simulations indicated that the population may continue to decline at >5% per year even after a mouse eradication.

Despite accounting for considerable uncertainty, it was very unlikely (<10%) that the MacGillivray’s prion population would become extremely vulnerable to extinction if mice were successfully eradicated in 2021 (Fig. 2). By contrast, without mouse eradication, and therefore persistent low productivity, there was a 31.5% probability that by 2057 the prion population would drop below 250 breeding pairs and thus become extremely vulnerable to extinction (Fig. 2).

**Figure 2.**
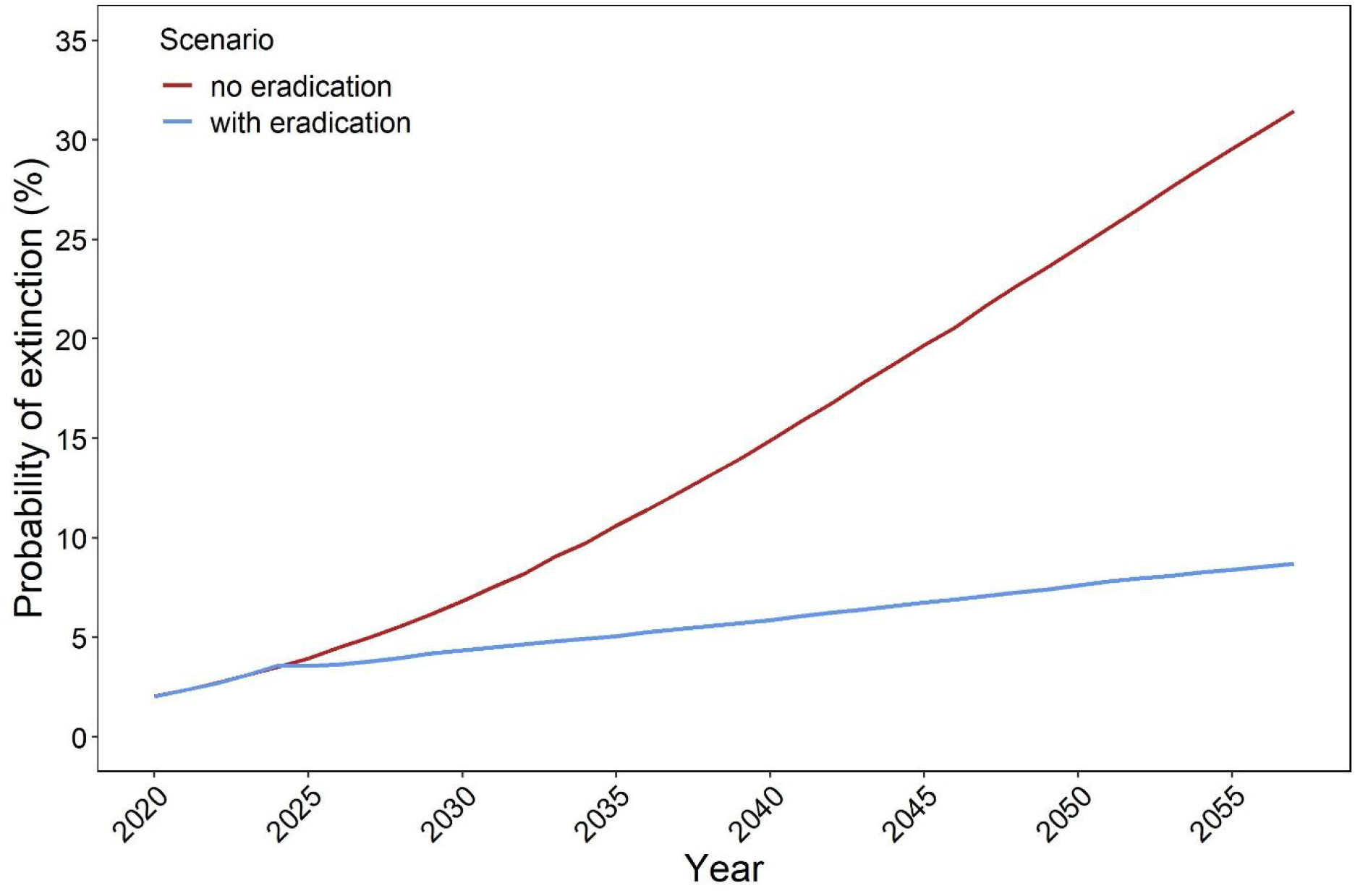
Estimated probability of becoming extremely vulnerable to extinction (<250 breeding pairs) for MacGillivray’s prions on Gough Island with and without a mouse eradication planned in 2021, based on 100,000 stochastic simulations in a population model using the demographic parameters presented in Table 1.

## Discussion

We show that MacGillivray’s prions on Gough Island continue to suffer heavy chick predation (Dilley et al. 2015) and that their population is likely declining rapidly despite reasonably high annual adult survival. Our demographic parameter estimates are consistent with the loss of more than 2 million breeding pairs on Gough between 1956 and 2000 and indicate that the species could become extremely vulnerable to extinction within the next three generations without mouse eradication. A successful mouse eradication would immediately improve annual productivity and this may be sufficient to stabilise the MacGillivray’s prion population on Gough. However, population recovery would likely be slow and centuries might be required to restore the population to similar abundance as estimated in 1956 (Jones 2010).

Only 0-2% of MacGillivray’s prion pairs fledged a chick in five out of six years of monitoring. This poor productivity is in stark contrast to the mean reproductive success of closely related prion species in predator-free environments (0.38 – 0.83; Van Rensburg and Bester 1988; Liddle 1994; Quillfeldt et al. 2003; Nevoux and Barbraud 2006; Quillfeldt et al. 2007), which our study colony only achieved in a single year. The winter preceding the 2016 breeding season had slightly more rainfall than the other five years of our data series (South African Weather Service, unpublished data), which may have resulted in a lower mouse population and reduced mouse predation on MacGillivray’s prion in that year (McClelland et al. 2018). The occasional occurrence of years with normal breeding success may have allowed the population to persist for the last decades, but ameliorating conditions under climate change may reduce the frequency of these ‘good’ breeding years (McClelland et al. 2018).

However, the infrequent occurrence of good breeding years cannot have persisted on Gough Island since 1956, as the predicted population growth rate (0.908, Table 1) given such a low frequency of good breeding years would have resulted in a much faster population decline than that observed between 1956 and 2000. We therefore cautiously inferred that good breeding years in 1956 likely occurred at much greater frequency (0.91, Table 1), and that this frequency declined on average at a rate of 0.013 per year (Table 1), which would be consistent with mouse population increases expected under climate change (McClelland et al. 2018). However, no data exist to support these assumptions, and it is possible that productivity declined more abruptly, or from a much higher original level. Regardless of the original productivity on Gough before mouse predation became widespread, it is likely that mouse predation will continue at or above the levels observed between 2014 and 2020, leading to breeding success well below the level required for population stability. We predict that under current conditions the MacGillivray’s prion population on Gough Island will become extremely vulnerable to extinction in the next three decades.

Our model projections are sensitive to the estimates of adult and juvenile survival, and the uncertainty in these parameters is reflected in the uncertainty of our future projections. Our adult survival estimate from ringing data indicates that MacGillivray’s prions on Gough Island have a reasonably high adult survival for a small to medium sized burrowing petrel, similar to adults on a predator-free islet in the Indian Ocean (Jiguet et al. 2007). However, our estimates for juvenile survival were taken from a declining shearwater species (Oro et al. 2004), and it is possible that in the absence of major anthropogenic threats at sea MacGillivray’s Prions may actually have higher juvenile survival similar to small gadfly petrels in the Pacific (Jones et al. 2011). Despite this uncertainty over annual survival probability of adults and juveniles, an improvement in breeding success following mouse eradication appears to be the only feasible conservation management intervention that could change the downward trajectory of the MacGillivray’s prion population on Gough Island.

The fate of MacGillivray’s prions is likely representative for several other small burrow-nesting species (e.g. storm petrels, diving petrels etc.) on Gough that are likely to be severely affected by mice (Ryan 2010; Cuthbert et al. 2013; Caravaggi et al. 2018), but for which no demographic data exist. Given the historical abundance of a number of species that are now locally scarce (Swales 1965; Ryan 2010; Dilley et al. 2015), it is highly likely that other burrow-nesting species such as broad-billed prion, Subantarctic shearwater *Puffinus elegans,* common diving petrel *Pelecanoides urinatrix* and three species of storm petrels (Oceanitidae) that breed on Gough Island have experienced similar population collapses as the one we show for MacGillivray’s prion. Caravaggi et al. (2018) found that the breeding success on Gough Island depends not only on body size and nesting substrate, but also on timing of breeding, with winter breeding species being more affected by mice.

MacGillivray’s prions breed in summer, hence species of similar or smaller size that breed at other times of the year – for example *Fregetta* spp. storm petrels, common diving petrel and broad-billed prion – could suffer even more severe predation than MacGillivray’s prion (Caravaggi et al. 2018). Thus, many small petrel species may become locally extinct on Gough should such high-intensity mouse predation continue. Therefore, the long-term persistence of several seabird species on Gough Island is dependent on the eradication of mice so that recruitment can be restored to levels required for population stability.

Invasive mammal eradication is an effective conservation intervention that usually leads to growth of seabird populations (Jones et al. 2016; Brooke et al. 2018; Holmes et al. 2019), which is reassuring for funders and practitioners who implement eradication operations to restore seabird breeding islands. Our simulations suggested that even when considering mice-free conditions and high productivity, the MacGillivray’s prion population might require centuries to recover to initial population levels (Table 1, Fig. 1), or may continue to decline if actual survival is at the lower margin of the range of survival values that we included in our simulations. Although the precision of our quantitative projections of population sizes is low, our comparison of alternative management strategies clearly shows that mouse eradication is urgently needed to prevent ongoing declines of Gough Island’s small petrel species.

## Acknowledgements

We thank all field staff from 2014 to 2018 for carrying out the monitoring at Prion Cave. Logistical support was provided by the South African Department of Environmental Affairs, the University of Cape Town and Royal Society for the Protection of Birds. The Tristan da Cunha Administrator, Island Council and Conservation Department provided permission to work on Gough Island. J. Ramos and two anonymous reviewers provided constructive comments on a previous version of this manuscript.

## Compliance with ethical standards

### Ethical approval

The Science Animal Ethics Committee (SFAEC) of the University of Cape Town approved the research protocols for this study (protocol number 2014/V10/PR) as part of a larger project entitled, “Population dynamics and conservation of Southern Ocean albatrosses and petrels”. Field procedures were approved by the Animal Ethics Committees of the Royal Society for the Protection of Birds.

## SUPPLEMENTARY MATERIAL

**Table S1:** Model selection table of four different parameterisations of the survival component of our population model, including either constant (.) or annually varying (time) probabilities of survival (phi) and recapture (p). DIC is the deviance information criterion, AIC_*c*_ is the Akaike Information Criterion adjusted for small sample size – both criteria indicate that the model formulation with constant survival and annually varying recapture probability is most parsimonious.

| Model | Deviance | N<br>parameters | DIC | ΔDIC | $AIC_c$ | $\Delta$ $AIC_c$ |
| --- | --- | --- | --- | --- | --- | --- |
| $\phi(.)$ $p(\text{time})$ | 68.90 | 6 | 73.68 | 0.00 | 81.48 | 0.00 |
| $\phi(\text{time})$ $p(\text{time})$ | 72.04 | 11 | 78.75 | 5.07 | 95.94 | 14.46 |
| $\phi(\text{time})$ $p(.)$ | 184.95 | 6 | 187.86 | 114.18 | 197.53 | 116.05 |
| $\phi(.)$ $p(.)$ | 208.06 | 2 | 210.04 | 136.36 | 212.14 | 130.65 |

**Fig. 1:**
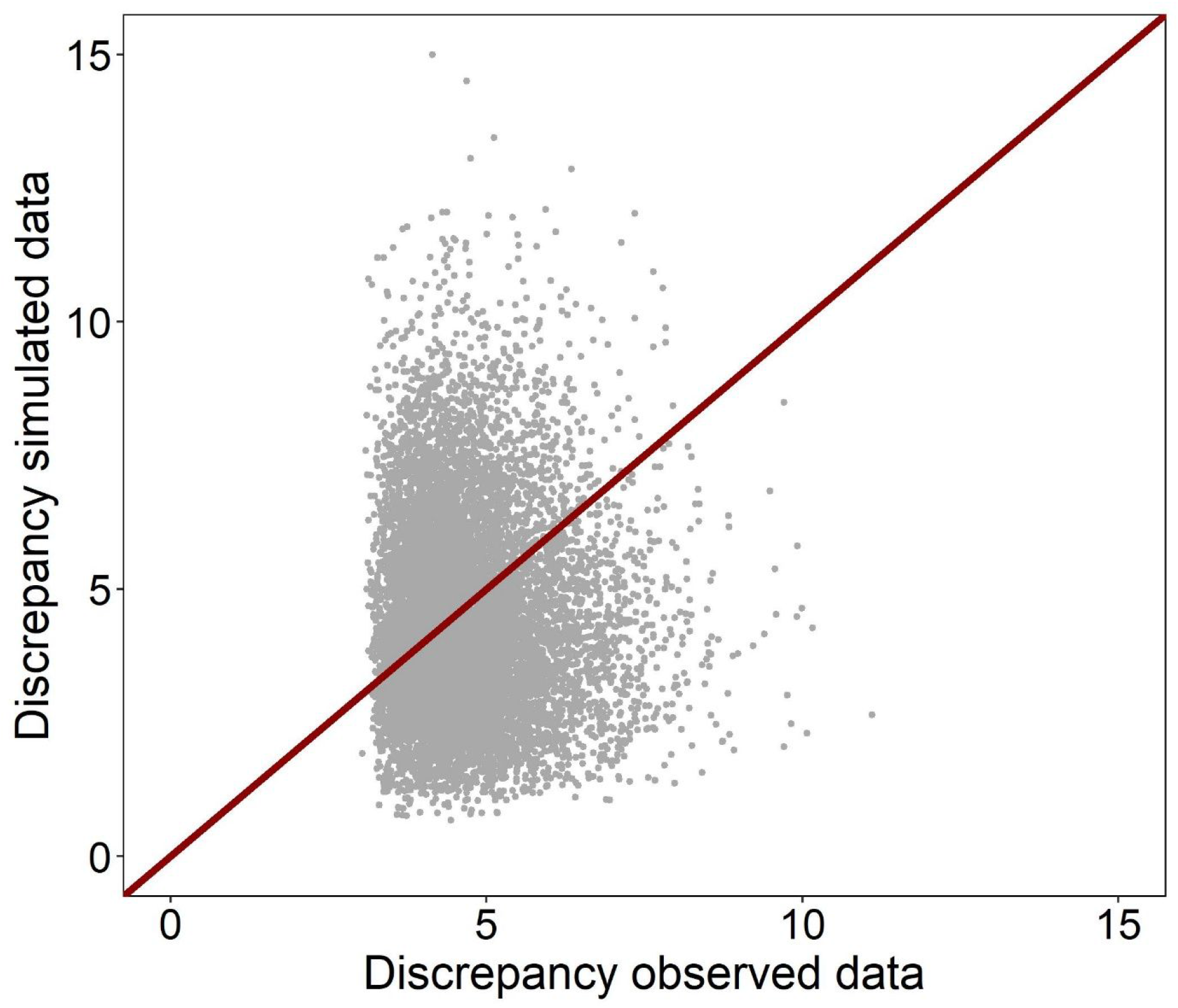
Posterior predictive check of model fit by a scatter plot of the discrepancy measure for simulated versus observed data in our survival model including all individually marked birds. The Bayesian p-value is the proportion of points above the 1:1 diagonal (0.42).

## References

Abadi, F., Gimenez, O., Ullrich, B., Arlettaz, R., Schaub, M. (2010). Estimation of immigration rate using integrated population models. J. Appl. Ecol. 47, 393–400.

Bird, J. P., Martin, R., Akçakaya, H. R., Gilroy, J., Burfield, I. J., Garnett, S. T., Symes, A., Taylor, J., Şekercioğlu, Ç. H., Butchart, S. H. M. (2020). Generation lengths of the world’s birds and their implications for extinction risk. Conservation Biology, doi:10.1111/cobi.13486.

Birdlife-International (2020). Species factsheet: Pachyptila macgillivrayi. Downloaded from http://www.birdlife.org on 28 July 2020,

Bretagnolle, V., Zotier, R., Jouventin, P. (1990). Comparative population biology of four prions (genus *Pachyptila*) from the Indian Ocean and consequences for their taxonomic status. The Auk 107, 305–316.

Brooke, M. (2004). Albatrosses and petrels across the world. Oxford University Press: Oxford.

Brooke, M. d. L., Bonnaud, E., Dilley, B., Flint, E., Holmes, N., Jones, H., Provost, P., Rocamora, G., Ryan, P., Surman, C. (2018). Seabird population changes following mammal eradications on islands. Animal Conservation 21, 3–12.

Brooke, M. d. L., O’Connell, T. C., Wingate, D., Madeiros, J., Hilton, G. M., Ratcliffe, N. (2010). Potential for rat predation to cause decline of the globally threatened Henderson petrel *Pterodroma atrata*: evidence from the field, stable isotopes and population modelling. Endang. Spec. Res. 11, 47–59.

Brooks, S., King, R., Morgan, B. (2004). A Bayesian approach to combining animal abundance and demographic data. Animal Biodiversity and Conservation 27, 515–529.

Broome, K., Brown, D., Brown, K., Murphy, E., Birmingham, C., Golding, C., Corson, P., Cox, A., Griffiths, R. (2019). House mice on islands: management and lessons from New Zealand. In: Veitch, C. R., Clout, M. N., Martin, A. R., Russell, J. C. (eds) Island invasives: scaling up to meet the challenge. IUCN: Dundee, pp 100–107.

Burnham, K. P., Anderson, D. R. (2002). Model selection and multimodel inference. A practical information-theoretic approach. 2nd edn. Springer: New York.

Caravaggi, A., Cuthbert, R. J., Ryan, P. G., Cooper, J., Bond, A. L. (2018). The impacts of introduced House Mice on the breeding success of nesting seabirds on Gough Island. Ibis 161, 648–661.

Conn, P. B., Johnson, D. S., Williams, P. J., Melin, S. R., Hooten, M. B. (2018). A guide to Bayesian model checking for ecologists. Ecol. Monogr. 88, 526–542.

Cuthbert, R. (2004). Breeding biology of the Atlantic Petrel, *Pterodroma incerta*, and a population estimate of this and other burrowing petrels on Gough Island, South Atlantic Ocean. Emu 104, 221–228.

Cuthbert, R., Hilton, G. (2004). Introduced house mice *Mus musculus:* a significant predator of threatened and endemic birds on Gough Island, South Atlantic Ocean? Biological Conservation 117, 483–489.

Cuthbert, R. J., Louw, H., Lurling, J., Parker, G., Rexer-Huber, K., Sommer, E., Visser, P., Ryan, P. G. (2013). Low burrow occupancy and breeding success of burrowing petrels at Gough Island: a consequence of mouse predation. Bird Conservation International 23, 113–124.

Davies, D., Dilley, B., Bond, A., Cuthbert, R., Ryan, P. (2015). Trends and tactics of mouse predation on Tristan Albatross *Diomedea dabbenena* chicks at Gough Island, South Atlantic Ocean. Avian Conservation and Ecology 10,

Dias, M. P., Martin, R., Pearmain, E. J., Burfield, I. J., Small, C., Phillips, R. A., Yates, O., Lascelles, B., Borboroglu, P. G., Croxall, J. P. (2019). Threats to seabirds: a global assessment. Biological Conservation 237, 525–537.

Dilley, B. J., Davies, D., Bond, A. L., Ryan, P. G. (2015). Effects of mouse predation on burrowing petrel chicks at Gough Island. Antarct. Sci. 27, 543–553.

Dilley, B. J., Schoombie, S., Schoombie, J., Ryan, P. G. (2016). ‘Scalping’of albatross fledglings by introduced mice spreads rapidly at Marion Island. Antarctic Science 28, 73–80.

Dilley, B. J., Schoombie, S., Stevens, K., Davies, D., Perold, V., Osborne, A., Schoombie, J., Brink, C. W., Carpenter-Kling, T., Ryan, P. G. (2018). Mouse predation affects breeding success of burrow-nesting petrels at sub-Antarctic Marion Island. Antarctic Science 30, 93–104.

Faulquier, L., Fontaine, R., Vidal, E., Salamolard, M., Le Corre, M. (2009). Feral cats Felis catus threaten the endangered endemic Barau’s petrel Pterodroma baraui at Reunion Island (Western Indian Ocean). Waterbirds 32, 330–336.

Frankham, R., Bradshaw, C. J. A., Brook, B. W. (2014). Genetics in conservation management: Revised recommendations for the 50/500 rules, Red List criteria and population viability analyses. Biol. Conserv. 170, 56–63.

Genovart, M., Pradel, R. (2019). Transience effect in capture-recapture studies: The importance of its biological meaning. PLOS ONE 14, e0222241.

Holmes, N. D., Spatz, D. R., Oppel, S., Tershy, B., Croll, D. A., Keitt, B., Genovesi, P., Burfield, I. J., Will, D. J., Bond, A. L. (2019). Globally important islands where eradicating invasive mammals will benefit highly threatened vertebrates. PLoS One 14, e0212128.

Horn, S., Greene, T., Elliott, G. (2019). Eradication of mice from Antipodes Island, New Zealand. In Island invasives: scaling up to meet the challenge: 131-136. Veitch, C. R., Clout, M. N., Martin, A. R., Russell, J. C., West, C. J. (Eds.). IUCN: Gland, Switzerland.

Jiguet, F., Robert, A., Micol, T., Barbraud, C. (2007). Quantifying stochastic and deterministic threats to island seabirds: last endemic prions face extinction from falcon peregrinations. Animal Conservation 10, 245–253.

Jones, C., Clifford, H., Fletcher, D., Cuming, P., Lyver, P. (2011). Survival and age-at-first-return estimates for grey-faced petrels (Pterodroma macroptera gouldi) breeding on Mauao and Motuotau Island in the Bay of Plenty, New Zealand. Notornis 58, 71–80.

Jones, C. W., Phillips, R. A., Grecian, W. J., Ryan, P. G. (2020). Ecological segregation of two superabundant, morphologically similar, sister seabird taxa breeding in sympatry. Marine Biology 167, 1–16.

Jones, C. W., Risi, M. M., Cleeland, J., Ryan, P. G. (2019). First evidence of mouse attacks on adult albatrosses and petrels breeding on sub-Antarctic Marion and Gough Islands. Polar Biol. 42, 619–623.

Jones, C. W. P. (2018). Comparative ecology of Pachyptila species breeding sympatrically at Gough Island. MSc thesis, University of Cape Town,

Jones, H. P. (2010). Seabird islands take mere decades to recover following rat eradication. Ecol. Appl. 20, 2075–2080.

Jones, H. P., Holmes, N. D., Butchart, S. H., Tershy, B. R., Kappes, P. J., Corkery, I., Aguirre-Muñoz, A., Armstrong, D. P., Bonnaud, E., Burbidge, A. A. (2016). Invasive mammal eradication on islands results in substantial conservation gains. Proceedings of the National Academy of Sciences 113, 4033–4038.

Keitt, B. S., Wilcox, C., Tershy, B. R., Croll, D. A., Donlan, C. J. (2002). The effect of feral cats on the population viability of black-vented shearwaters *(Puffinus opisthomelas)* on Natividad Island, Mexico. Animal Conservation 5, 217–223.

Kellner, K. F. (2016). jagsUI: A wrapper around rjags to streamline JAGS analyses. Available at: https://CRAN.R-project.org/package=jagsUI, accessed

Kéry, M., Schaub, M. (2012). Bayesian population analysis using WinBUGS. Academic Press: Oxford, UK.

Lavers, J. L., Wilcox, C., Donlan, C. J. (2010). Bird demographic responses to predator removal programs. Biological Invasions 12, 3839–3859.

Liddle, G. (1994). Interannual variation in the breeding biology of the Antarctic prion *Pachyptila desolata* at Bird Island, South Georgia. Journal of Zoology 234, 125–139.

McClelland, G. T. W., Altwegg, R., van Aarde, R. J., Ferreira, S., Burger, A. E., Chown, S. L. (2018). Climate change leads to increasing population density and impacts of a key island invader. Ecol. Appl. 28, 212–224.

McClelland, P. (2019). Operational plan for the eradication of House Mice from Gough Island. In. Royal Society for the Protection of Birds: Sandy, UK

Nevoux, M., Barbraud, C. (2006). Relationships between sea ice concentration, sea surface temperature and demographic traits of thin-billed prions. Polar biology 29, 445–453.

Oro, D., Aguilar, J. S., Igual, J. M., Louzao, M. (2004). Modelling demography and extinction risk in the endangered Balearic shearwater. Biol. Conserv. 116, 93–102.

Plard, F., Fay, R., Kéry, M., Cohas, A., Schaub, M. (2019). Integrated population models: powerful methods to embed individual processes in population dynamics models. Ecology 100, e02715.

Plummer, M. (2012). JAGS Available at: http://sourceforge.net/projects/mcmc-jags/files/Manuals/, accessed 12 Aug 2013.

Pradel, R., Hines, J. E., Lebreton, J. D., Nichols, J. D. (1997). Capture-recapture survival models taking account of transients. Biometrics 53, 60–72.

Quillfeldt, P., Masello, J. F., Strange, I. J. (2003). Breeding biology of the thin-billed prion *Pachyptila belcheri* at New Island, Falkland Islands: egg desertion, breeding success and chick provisioning in the poor season 2002/2003. Polar Biology 26, 746–752.

Quillfeldt, P., Strange, I. J., Masello, J. F. (2007). Sea surface temperatures and behavioural buffering capacity in thin-billed prions *Pachyptila belcheri:* breeding success, provisioning and chick begging. Journal of Avian Biology 38, 298–308.

R-Core-Team (2019). R Project for Statistical Computing. R Foundation for Statistical Computing. Available online: www.R-project.org: Vienna.

Rayner, M. J., Hauber, M. E., Imber, M. J., Stamp, R. K., Clout, M. N. (2007). Spatial heterogeneity of mesopredator release within an oceanic island system. Proceedings of the National Academy of Sciences 104, 20862–20865.

Russell, J. C., Peace, J. E., Houghton, M. J., Bury, S. J., Bodey, T. W. (2020). Systematic prey preference by introduced mice exhausts the ecosystem on Antipodes Island. Biological Invasions 22, 1265–1278.

Ryan, P. (2010). Tipping point: mice eroding Gough’s seabirds. Africa Birds & Birding 15, 13.

Ryan, P. G., Bourgeois, K., Dromzée, S., Dilley, B. J. (2014). The occurrence of two bill morphs of prions *Pachyptila vittata* on Gough Island. Polar biology 37, 727–735.

Schaub, M., Abadi, F. (2011). Integrated population models: a novel analysis framework for deeper insights into population dynamics. J. Orn. 152, 227–237.

Schaub, M., Gimenez, O., Sierro, A., Arlettaz, R. (2007). Use of integrated modeling to enhance estimates of population dynamics obtained from limited data. Conserv. Biol. 21, 945–955.

Swales, M. (1965). The sea-birds of Gough Island. Ibis 107, 215–229.

Tollu, B. (1984). La Quille (île Saint Paul, Océan Indien), sanctuaire de populations relictes. Oiseau et la Revue Francaise d’Ornithologie 54, 79–85.

Van Rensburg, P., Bester, M. (1988). The effect of cat Felis catus predation on three breeding Procellariidae species on Marion Island. South African Journal of Zoology 23, 301–305.

Wade, P. R. (2000). Bayesian methods in conservation biology. Conserv. Biol. 14, 1308–1316.

Wanless, R. M., Angel, A., Cuthbert, R. J., Hilton, G. M., Ryan, P. G. (2007). Can predation by invasive mice drive seabird extinctions? Biology Letters 3, 241–244.

Wanless, R. M., Ryan, P. G., Altwegg, R., Angel, A., Cooper, J., Cuthbert, R., Hilton, G. M. (2009). From both sides: Dire demographic consequences of carnivorous mice and longlining for the Critically Endangered Tristan albatrosses on Gough Island. Biological Conservation 142, 1710–1718.

Warham, J. (1990). The petrels: their ecology and breeding systems. A&C Black: London.

Worthy, T. H., Jouventin, P. (1999). The fossil avifauna of Amsterdam Island, Indian Ocean. Smithsonian contributions to paleobiology 89, 39–65.

Zipkin, E. F., Saunders, S. P. (2018). Synthesizing multiple data types for biological conservation using integrated population models. Biol. Conserv. 217, 240–250.

